# T-FINDER: A highly sensitive, pan-HLA platform for functional T cell receptor and ligand discovery

**DOI:** 10.1101/2023.05.16.540992

**Authors:** Theresa Schmid, Miray Cetin, Veronica Pinamonti, Ana Mellado Fuentes, Kristina Kromer, Taga Lerner, Jing Zhang, Tamara Boschert, Yonatan Herzig, Christopher Ehlert, Laura Fisch, Valeriia Dragan, Arlette Kouwenhoven, Bertrand Van Schoubroeck, Hans Wils, Carl Van Hove, Michael Platten, Edward Green, Frederik Stevenaert, Nathan J. Felix, John M. Lindner

## Abstract

Effective, unbiased, high-throughput methods to functionally identify both class II and class I HLA-presented T cell epitopes and their cognate T cell receptors (TCRs) are essential for and prerequisite to diagnostic and therapeutic applications, yet remain underdeveloped. Addressing this bottleneck, we established T-FINDER (T cell Functional Identification and (Neo)-antigen Discovery of Epitopes and Receptors), a platform that rapidly deconvolutes CD4 and CD8 TCR reactivities to targets physiologically processed and presented by an individual’s unmanipulated, complete HLA haplotype. By using a highly sensitive TCR signaling reporter capable of detecting even low-affinity TCR:ligand interactions, T-FINDER not only robustly identifies unknown peptide:HLA ligands from complex antigen libraries, but also rapidly screens and functionally validates the specificity of complex TCR libraries against known or predicted targets. To demonstrate its pan-HLA presentation capacity, we apply the platform to multiple TCR-based applications, including glioma, celiac disease, and rheumatoid arthritis, providing unique biological insights and showcasing T-FINDER’s potency and versatility.

## Introduction

The molecular interaction between T cell receptors (TCRs) and their cognate ligands instructs many functions and dysfunctions of the adaptive immune system. While CD8+ T cell responses directly effect the killing of infected or transformed somatic cells, CD4+ T cells interact with professional antigen-presenting cells (APCs) to orchestrate a range of immune responses, including inducing a proinflammatory state, promoting and defining antibody production, and driving cytotoxicity. Conversely, TCR engagement can induce immunosuppression via the action of regulatory T cells. Each T cell expresses a somatically rearranged, clonally unique TCR, and next-generation sequencing can generate repertoire data from virtually any patient tissue or *ex vivo* manipulated T cell population, providing lists of “TCRs of interest” based on phenotype and/or level of clonal expansion^1^, but no specificity information. Understanding the cognate specificity of disease-relevant TCRs can provide insights into etiopathology and guide therapeutic strategies. However, functional approaches to deconvolute *de novo* TCR-ligand interactions and validate predicted ligands remain technologically challenging. Existing approaches pairing TCR sequences with their cognate ligands include cell-based reporter systems, but are mostly applied to CD8 TCRs and class I HLA-presented antigens^2, 3^. Several more recently established platforms are capable of CD4 TCR deconvolution, but their general utility, scalability, and capacity for true *de novo* epitope discovery remain unclear^4, 5^. Similarly, *in silico* approaches for predicting the identity and HLA restriction of a TCR ligand, while offering promising throughput and efficiency, are limited by the quality of their training datasets, and generalize poorly to novel epitopes that do not share structural similarity to the training data^6, 7^. As a result, many TCR clones and their functional ligands associated with disease pathogenesis remain unknown.

To identify functional TCR ligands and overcome the aforementioned limitations, we designed a highly sensitive TCR deconvolution platform to accomplish the following: maintain the physiological TCR:target interaction without molecular engineering, be agnostic to predicted epitopes and presenting alleles, work for both class II and class I-presented peptides, and be independent from mass spectrometry, tetramer labeling, and other ligand/TCR prediction steps. This system, capable of bias-free *de novo* ligand identification given a TCR of interest, is sufficiently powerful to provide rapid TCR deconvolution using complex antigen libraries, as well as facile validation of predicted TCR:ligand interactions. After establishing and benchmarking our platform, we performed functional ligand deconvolution screens across a series of infectious disease, autoimmune, and oncologic settings, demonstrating the flexibility and potency of the system. Based on its ability to physiologically identify and intrinsically validate TCRs and their epitopes, we named this platform T-FINDER (T cell Functional Identification and (Neo)-antigen Discovery of Epitopes and Receptors).

## Materials and Methods

### Cell lines

T cell reporter lines with and without TCRs were cultured in GlutaMAX^TM^ supplemented RPMI 1640 medium further modified with 10% heat-inactivated FBS, 100 U mL^-1^penicillin and 100 µg mL^-1^ streptomycin, 1 mM sodium pyruvate, and 55 μM β-mercaptoethanol. TCR signaling reporter lines were additionally cultured with 10 µg mL^-1^ blasticidin and TCR-transgenic lines were selected using 1 µg mL^-1^ puromycin. The human lymphoblastoid line BOLETH was obtained from Sigma (#88052031-1VL). B-cell lines were cultured in GlutaMAX^TM^ supplemented RPMI 1640 medium further modified with 10% FBS, 100 U mL^-1^ penicillin and 100 µg mL^-1^ streptomycin, 1 mM sodium pyruvate, 1x MEM Non-Essential Amino Acids Solution, and 55 μM β-mercaptoethanol. HEK293T cells were cultured in high-glucose GlutaMAX^TM^ supplemented DMEM containing 10% heat-inactivated FBS, 100 U mL^-1^ penicillin and 100 µg mL^-1^ streptomycin. DPBS without calcium or magnesium and 0.05% Trypsin-EDTA was used for passaging. All human cell lines were cultured at 37°C with 5% CO_2_ in a humidified atmosphere.

### Primary cell culture and B-LCL immortalization

For Epstein-Barr virus-induced immortalization of donor-derived B cells, PBMCs were isolated from buffy coats using 50 mL Leucosep^TM^ centrifuge tubes, Ficoll Paque Plus, and ACK erythrocyte lysis buffer. B cells were extracted by negative selection with the EasySep™ Human B Cell Isolation Kit (StemCell Technologies, #17954). EBV-containing supernatants from B95-8 cells were added to purified B cells supported by 30 µg mL^-1^ holo-transferrin and 2.5 µg mL^-1^ CpG (ODN2006). Established lines were haplotyped by DKMS Life Science. The primary tumor cell line used for antigen presentation to CAR1 and CAR2 was kindly provided by Janssen.

### Development of the reporter construct and T cell reporter line

Synthetic TCR signaling-response promoter sequences (Supplemental Table 1) were cloned 5’- of the reporter cassette in a Sleeping Beauty transposon backbone. This cassette, protected by core insulator sequences, harbors an HA-tagged αCD19 (B43^8^) scFv sequence followed by a furin cleavage site, T2A viral peptide, and superfolder GFP^9^. The 3’-UTR of human IgG1 was used to increase transcript stability and provide a poly-adenylation signal. Following this inducible cassette and a core insulator sequence, a constitutively active PGK promoter drives the expression of CD28 and a blasticidin resistance gene. A separate plasmid containing the transposase was co-transfected (1:5). All DNA sequences were synthesized by Twist Bioscience. To generate the T cell reporter line, 2.5x10^6^ TCR-deficient Jurkat or SKW-3 cells were electroporated with plasmid DNA using the Neon Transfection system (Thermo Fisher). Single-cell clones were expanded under selection (8 or 10 µg mL^-1^ blasticidin) for two weeks before measuring TCR-inducible reporter activity.

### Design and construction of TCR reporter cells

T cell receptors were generated by custom synthesis (Twist Bioscience) in a 3^rd^ generation pLVX-EF1α-IRES-Puro lentiviral vector (Takara/Clontech). Following the TransIT-Virusgen Transfection (Mirus, #MIR6700) protocol, HEK293T cells were used with VSV-G envelope expressing plasmid pMD2.G and packaging plasmid psPAX2. Jurkat and SKW-3 reporter cells were transduced with filtered viral supernatant and 1 µg mL^-1^ polybrene 48h after transfection. Viral supernatant was replaced with fresh medium after 24h and selected with 1 µg mL^-1^ puromycin 96h after transduction. CAR1 and CAR2 were provided as filtered viral supernatants by Janssen and processed as described above.

### T cell receptor sequences

The glioma patient (ID07) received H3K27M-vac after providing written, signed informed consent at the University Hospital of Mannheim with treatment and sample use approved by the institutional review board. Other sequences were reconstituted as full-length TCRs from published gene segment usage and CDR3 information or identified following TCR sequencing of pseudonymized cellular material upon provision of informed consent for experimental purposes.

### Design, cloning, and transgenesis of genetically templated antigens

Epitope-containing minigene sequences were generated by custom synthesis (Twist Bioscience). These sequences contained an open reading frame (up to 400 aa) of either a full-length gene of interest or fragment thereof. Smaller fragments were amplified by PCR from minigenes with custom primers (Integrated DNA Technologies) using Q5 high-fidelity DNA Polymerase. Each construct was flanked by unique SpeI and BamHI sites to enable efficient cloning into a pLVX-EF1α-IRES-Puro lentiviral vector, which places the respective fragment upstream of an in-frame T2A-linked mTagBFP2^10^ sequence. Lentiviral transduction included an additional pre-incubation step of the B-LCLs with 4 nM BX-795 hydrochloride. Viral supernatant was replaced with fresh medium 24h after transduction. 72h after this medium change, the expression was verified by flow cytometric analysis of BFP and functional assays were performed.

### *In vitro* loaded peptides

Peptides were generated by custom peptide synthesis (Biomatik) or provided as peptide pools (JPT or Miltenyi BioTec, see SI for details). Peptides were resuspended at a stock concentration of 10 mg mL^-1^ in 10% DMSO and stored at -80°C. Peptide concentrations were adjusted in a dilution series with DMSO and an identical volume was added to B-LCL cells. The final concentrations of peptides used in co-cultures are shown in Figure 1 and Supplemental Figure 1.

**Figure 1.**
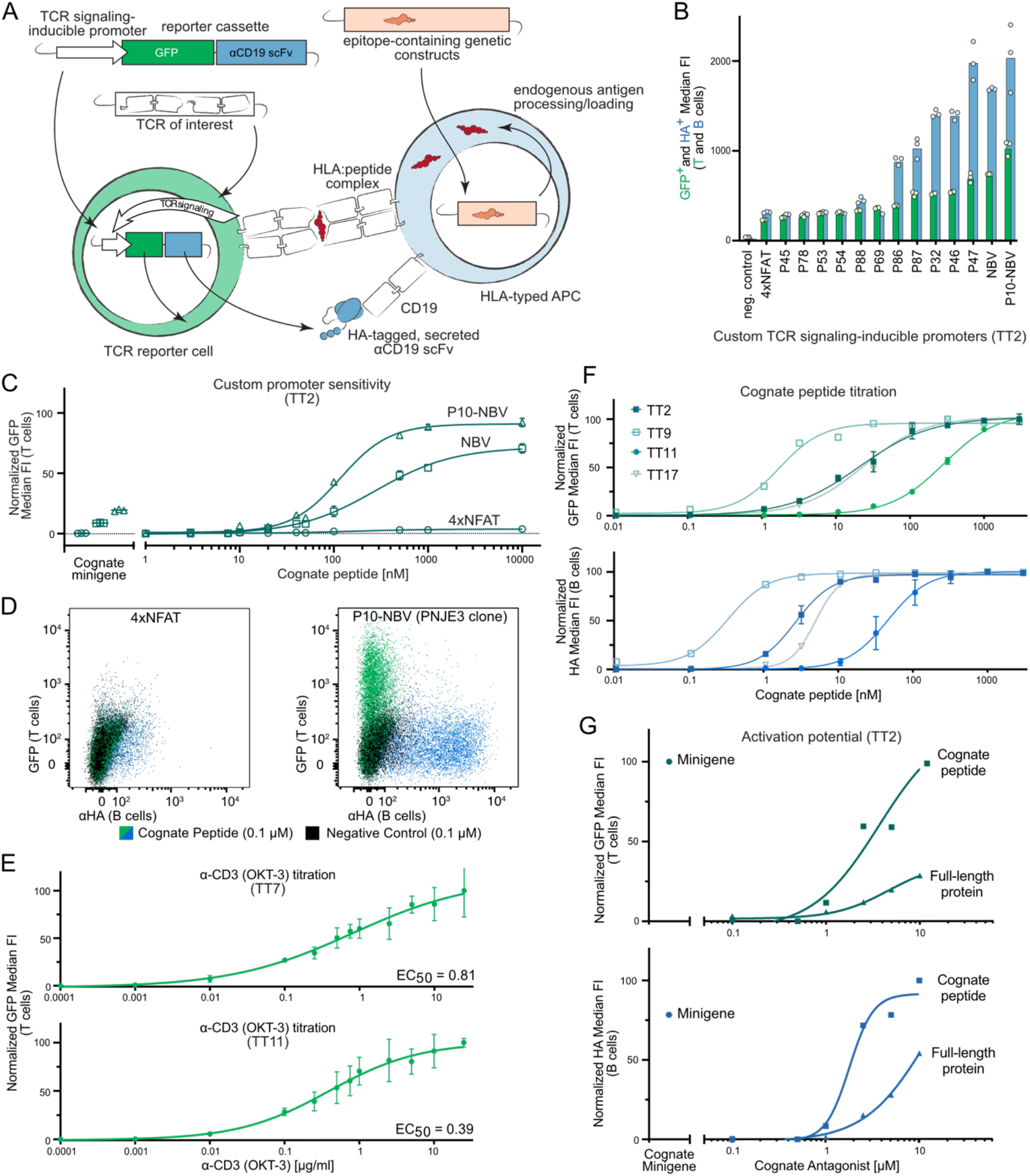
Developing and benchmarking a TCR activation reporter system. A) Schematic depicting construction of a TCR reporter cell line (left) transgenically modified with an inducible promoter and TCR of interest. Upon cognate ligand interaction, the reporter cell produces cytoplasmic GFP and an HA-labeled secreted antibody fragment specific for CD19, a B cell-specific surface protein. B cells (right) used as antigen-presenting cells are transduced with epitope containing minigenes or exogenously loaded with peptide epitopes. Upon reporter T cell activation, the B cells are labeled by the secreted αCD19 scFv. B) Median fluorescent GFP (green) and αHA (blue) signals upon cognate ligand interaction for TT2-transgenic Jurkat cells with bespoke TCR signaling-inducible promoters. Negative control, non-cognate interaction (from P10-NBV promoter as an example). C) Minigene and peptide dilution curves of a cognate epitope for P10-NBV, NBV, and 4xNFAT transposon-integrated inducible reporter cassettes (bulk T cell population prior to single-clone selection). D) Superimposed dot plots depicting T cell GFP expression (y-axis, green) and B-cell anti-CD19 scFv labeling (x-axis, blue) in a single cognate-target co-culture. Black dots, GFP fluorescence in a non-cognate control co-culture. Left, 4xNFAT promoter; right, P10-NBV promoter-transgenic single clone. E) TCR-independent titration curves for two benchmarking TCRs (TT7 and TT11) using an αCD3 antibody (OKT-3). F) Titration curves of PNJE3 activation of four benchmarking TCRs (TT2, TT9, TT11, and TT17) with cognate peptide epitopes from the *C. tetani* tetanospasmin protein. Upper plot, T cell GFP expression; lower plot, B-cell anti-CD19 scFv labeling. G) Activation of PNJE3 in co-culture with BOLETH cells following cognate peptide and full-length tetanus toxoid protein electropulsing, or transduction with an epitope-containing 400-amino acid minigene. Representative data from 2 independent titration experiments.

### Peptide and protein electroporation

Full-length inactivated tetanus toxoid (Uniprot ID P04958, 1315 aa) was kindly provided by Janssen, while the cognate 12-mer was generated by custom peptide synthesis (Biomatik). Both were resuspended at a stock concentration of 10 mg mL^-1^ in 10% DMSO and stored at -80°C. Per electroporation, 5x10^5^ B-LCL were washed with DPBS (without Ca^2+^ and Mg^2+^) and resuspended at 5x10^7^ cells mL^-1^ in 0.9% DPBS, adding full-length protein or peptide to the required final concentration. After electroporation (1300 V, 30 W, 1 P), cells were resuspended in complete, supplemented medium at 2x10^6^ cells mL^-1^. Electroporated cells recovered for 1h at 37 °C, 5% humidified CO_2_ before performing functional assays.

### T cell reporter stimulation and co-culture assays

Ligand-independent reporter cell activation was assessed by overnight stimulation with 6.25 µg mL^-1^ Ultra-LEAF™ Purified anti-human CD3 (OKT3, BioLegend, #317326). Cells were incubated in antibody-coated wells and TCR responses were measured by flow cytometric analysis of GFP expression after 16h. For co-culture experiments, 10^5^ reporter cells were seeded in U-bottom 96-well plates with fresh complete medium either 4 days after TCR transduction or following 2 additional weeks of selection (1 µg mL^-1^ puromycin). After 1h recovery in fully supplemented medium, antigen-presenting cells were added. Minigene-expressing APCs were co-cultured with reporter cells 4 days after transduction. In both approaches, antigen-presenting cells were combined with reporter cells at a 1:1 or 2:1 ratio in a final volume of 200 µL and incubated overnight (16h) at 37°C, 5% CO_2_ in a humidified atmosphere. Flow cytometry was used to assess TCR responses to HLA-presented epitopes through GFP expression and the B cell-bound HA-tagged αCD19 scFv fragment. T and B cells were identified using surface staining for CD2 and CD20, respectively. BFP was used to evaluate minigene translation efficiency.

### Flow cytometry analysis

The following antibodies from BioLegend were diluted in flow cytometry buffer (DPBS, 2% FBS, 2 mM EDTA): PerCP-Cy5.5-conjugated anti-human CD2 (RPA-2.10, 0.25 µg mL^-1^, #300216), PE-Cy7-conjugated anti-human CD3e (UCHT1, 0.75 µg mL^-1^, #300420), Pacific Blue or PE-Cy7-conjugated anti-human CD20 (2H7, 1 µg mL^-1^, #302320 or #302312), APC-Cy7 or PE-Cy7-conjugated anti-human CD69 (FN50, 0.5 µg mL^-1^, #310914 or #310912), and APC anti-HA.11 epitope tag (16B12, 0.5 µg mL^-1^, #901524). BV395-conjugated CD20 (2H7, 0.25 µg mL^-1^, #563782) was purchased from BDbiosciences. Propidium Iodide Ready Flow™ Reagent (Thermo Fisher, #R37169) was used as viability dye. GFP and BFP were additionally analyzed to identify TCR activation and minigene translation efficiency. Cells were washed before staining for 20 min at 4 °C in the dark. Samples were then washed and analyzed using a BD FACSAria Fusion or BD FACSAria II cell sorter, or in high throughput using a Sony ID7000 spectral analyzer. Data analysis was performed using FlowJo (10.6.1). For single-variable analysis, the percentage of positive cells was calculated as the integrated area under the histogram not overlapping with control samples using FlowJo’s population comparison (Overton) method.

### Design, co-culture, and imaging of multiplexed arrays

Following an algorithmically-generated pipetting scheme (code available on GitHub https://github.com/hits-ccc/Permutation), 5 independent minigenes were combined in each well of a low-DNA binding 96-well plate. 65 ng of each minigene resulted in a total of 0.325 µg of lentiviral vector in each 96-well. As described above, TransIT-Virusgen Transfection reagent was used, including 33 ng pMD2.G and 50 ng psPAX2 per well. 48h after transfection of virus-producing 293T cells, 5x10^4^ B-LCL cells were treated with BX-795 hydrochloride and 1 µg mL^-1^ polybrene for 15 min, before adding viral supernatant. For microscopy, B-LCL medium was exchanged for FluorBrite DMEM (Thermo Fisher, #A1896701) supplemented with 5% FBS, 100 U mL^-1^ penicillin and 100 µg mL^-1^, 1 mM sodium pyruvate, 1X MEM Non-Essential Amino Acids, and 55 μM β-mercaptoethanol 4 days after transduction. Transduced B-LCL cells of each 96 well were split into two wells of a clear-bottom 384-well plate. In an equal ratio, 2.5x10^4^ reporter cells selected for a TCR of interest were added to each 384 well. GFP expression was imaged following 7h of co-culture. Using the automatic scan function of the Nikon Eclipse Ti2 microscope, GFP images were recorded individually for each 384 well at 10x magnification (500ms). After removing the background, a macro for the software Fiji-ImageJ was applied to extract the GFP-positive area of each individually measured well. Z-scores were calculated as the number of standard deviations of each well above or below the mean using the distribution of all wells in a given experiment; geometric means from technical replicates were calculated to determine the signal threshold for positive wells. Minigenes shared among positive wells were identified as cognitive minigenes for the investigated TCR(s).

## Results

### Generating a TCR reporter system with high sensitivity and signal-to-noise ratio

To increase the discriminatory potential and flexibility of a TCR reporter, we designed a cassette that would, upon TCR signaling, provide both T cell-intrinsic and -extrinsic readouts. While the cytoplasmic production of GFP^9^ provides T cell fluorescence easily detectable by flow cytometry or microscopy, a secreted B-cell specific single-chain variable fragment (scFv) labels nearby APCs for direct identification, e.g., by flow cytometry (Figure 1A). This APC-labeling feature is highly advantageous as a secondary discriminating signal when working with weak TCRs and/or poorly presented epitopes, and crucial for high-throughput applications in which the stimulating APC (containing the genetic information of the cognate epitope) must be isolated from the activated reporter cells.

The dual-reporter cassette is preceded by an inducible promoter that potently responds to TCR engagement. One drawback of many TCR reporter systems is an insufficient signal in response to weak activation by low-affinity interactions. We generated a highly sensitive TCR reporter with a maximized signal-to-noise ratio by optimizing our synthetic promoter over several iterations, and confirmed its sensitivity using published TCR:pHLA combinations as well as internally generated TCRs as benchmarks (Supplemental Table 1-3). In particular for benchmarking purposes, we tested promoter strengths with a series of tetanus toxoid (TT) reactive TCRs (e.g., TT2, TT7, and TT11 shown in Figure 1). The epitopes of these TCRs had previously been mapped using peptide libraries, and their identities were later re-blinded to experimenters when validating deconvolution strategies.

Previously established promoter elements (e.g., 4 tandem copies of the NFAT binding sequences) did not yield adequate signal-to-noise ratios when placed into our reporter cassette (Figure 1B-D, Supplemental Figure 1A-B and D). Therefore, we designed a series of promoters (P1-P15) using re-assembled 60-bp genomic fragments from TCR and calcium signaling response genes^11, 12^ centered around NFAT binding sites, as well as fully synthetic combinations of transcription factor binding sequences. For the latter, in addition to NFAT consensus motifs, we included binding sites for NFκB, Fos/Jun (AP-1), and Oct1/2-Bob1 transcriptional activation complexes to maximally integrate TCR signals within the cell. These synthetic designs also accounted for binding motif spacing to minimize steric hindrance and sharing of transcriptional activators on the DNA^13^. While several such assembled promoters provided TCR-inducible activity, one synthetic construct (named ‘NBV’ after its NFAT-Bob1-Various structure) yielded a nearly 10-fold signal-to-noise ratio when integrated via a lentiviral transgene, increasing to nearly 50-fold when integrated via a genomically protected transposon (Supplemental Figure 1A). The design of a second series of promoters (P45-P90), incorporating fragments of the previously identified TCR-responsive elements, provided building blocks for ‘tunable’ TCR-inducible promoters with a range of signal strengths (Figure 1B). However, the greatest signal-to-noise ratio was observed when combining the synthetic NBV promoter with the highly inducible P10 genomic re-assembly fragment (P10-NBV, Figure 1B-C and Supplemental Figure 1B).

Following single-cell cloning and selection of a P10-NBV reporter clone (PNJE3) with the strongest signal-to-noise ratio (Figure 1D), we assessed the sensitivity of the resulting line to serial dilutions of anti-CD3 antibody (Figure 1E, Supplemental Figure 1C) and *in vitro* loaded peptides (Figure 1F). PNJE3 demonstrated nanomolar EC_50_ values with linear response ranges over several orders of magnitude combined with very low background signal. The reporter cassette itself is inducible in commonly used T cell lines including Jurkat (Figure 1B-G) and SKW-3 (Supplemental Figure 1D). Notably, the strong activation potential of the reporter does not show any background activity in the absence of a cognate ligand, indicating that the inducible promoter is tightly controlled, thereby reducing the likelihood of false positive signals. In an experiment testing 50 off-target TCRs, the T-FINDER reporter readouts were cleaner than CD69 surface expression in the absence of any presented cognate ligand, a typical proximal indicator of TCR signaling-mediated activation, suggesting that while CD69 levels reflect antigen-independent, tonic signaling of a given TCR, the T-FINDER reporter is only activated in response to complete T cell engagement by cognate ligands (Supplemental Figure 1E).

### Antigen presentation to reporter T cells

The TCR reporter system requires a cellular partner to present putative cognate peptide:HLA complexes. As T-FINDER is not restricted to a specific ligand-presenting cell, its compatibility with existing methods, such as tumor cell lines and engineered APCs^14^ provides a large degree of flexibility. However, when performing *de novo* target identification for a given donor, interrogating entire autologous HLA haplotypes simultaneously is most efficient. For this, we used Epstein-Barr virus-immortalized primary B lymphocytic cell lines (B-LCLs) as autologous APCs to maintain donor-specific aspects of antigen processing and presentation, most importantly the HLA system. Notably, B-LCLs were able to process full-length tetanus toxoid protein into peptide epitopes (Figure 1G), demonstrating that T-FINDER is compatible with physiological antigen processing pathways. Finally, we introduced genetically encoded antigen libraries as 400-amino acid protein fragments (“minigenes”). Delivery of minigene-expressed antigens to class II processing compartments was facilitated by the inclusion of a novel Processing Enhancing Sequence (PES) (Pinamonti et al, manuscript in preparation), which enabled the use of large minigene fragments that were otherwise inefficiently processed/presented to T cells. B-LCLs efficiently processed and presented minigene-encoded epitopes and activated cognate reporter T cells in the same range as B-LCL loaded with 10 nM peptide (Figure 1C, Supplemental Figure 1B). Notably, HLA zygosity did not impact the sensitivity of our reporter system; we obtained equivalent results using DR4-homozygous (BOLETH) and hemizygous B-LCLs (Supplemental Figure 1F), indicating that donor-derived B-cell lines, which recapitulate the entire complement of HLA alleles, can be used to identify TCR:epitope pairs.

### TCR ligand identification is rapid and robust across therapeutic indications

Designed to identify TCR ligands with no *a priori* information regarding peptide targets and presenting HLA alleles, T-FINDER’s sensitivity and signal strength are optimized to assess the functionality of TCRs and TCR-like receptors at many levels of scale, ranging from single-target validation assays to proteome-wide reactivity studies. These stringent performance requirements imply that less complex studies should be significantly faster and more informative than existing methodologies. To confirm this, we validated a series of TCR sequences for reactivity to peptides or minigene constructs up to 400 amino acids in length and containing their proposed and/or known cognate epitope(s) in disease-relevant settings, such as immuno-oncology, vaccinology, and infectious disease.

First, we tested a series of known TCRs specific for class I epitopes from cancer and infectious disease, including an anti-CMV TCR recently identified using a similar approach ^5^. All TCRs activated the PNJE3 reporter line (Figure 2A), confirming the platform’s sensitivity to class I presented minigene-encoded epitopes. We then screened clonally expanded CD4 TCRs from a diffuse midline glioma (DMG) patient administered a neo-epitope peptide vaccine^15^ to understand whether this treatment elicited a T cell driven immune response. We identified 20 unique TCR clonotypes reactive to a mutant histone 3 K27M vaccinating peptide (Figure 2B). While many TCR:ligand interactions generated robust activation signals, several combinations did not induce the PNJE3 line to produce large quantities of GFP, suggesting a weaker affinity and/or avidity of those TCR complexes to their cognate epitope(s). Therefore, we leveraged the integrated additional reporter label to optimize our quality control and analysis pipeline, ensuring that even weak responses are significantly above background for confident detection in high-throughput assays (Supplemental Figure 2). Using a series of single-allele HLA-deficient APC lines derived from the autologous B-LCL, we were also able to identify the class II alleles presenting the epitope to each TCR. In this patient, peptide-reactive TCRs restricted on both the HLA-DR and HLA-DQ loci were present (Supplemental Figure 3). These results highlight the sensitivity of the T-FINDER platform and breadth of the T cell response to this neo-epitope vaccine (for additional donors and more detailed analyses, see Boschert, Kromer, Lerner et al 2023 BioRxiv^16^ and Grassl N et al, in revision).

**Figure 2.**
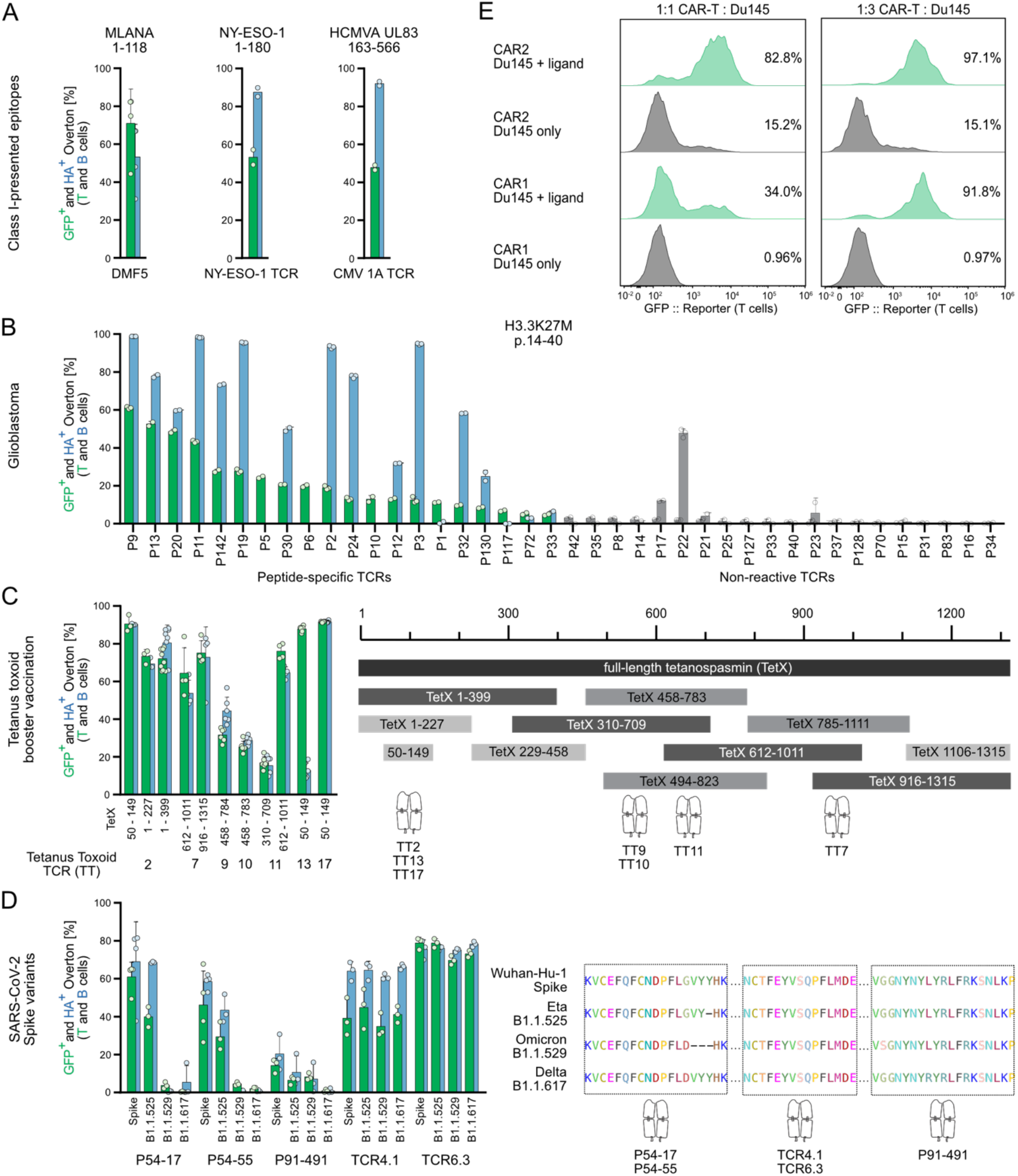
Functional assessment of T cell:ligand interactions. A) PNJE3 activation by class I-restricted TCRs co-cultured with HLA-matched, cognate minigene-transduced B-LCLs. B) Rapid screening of putative neo-epitope reactive TCRs in H3K27M diffuse midline glioma (DMG). TCR IDs indicate their clonal frequency following *ex vivo* epitope-specific enrichment. Clonotypes are plotted in order of descending GFP signal following co-culture with a peptide-loaded autologous B-cell line. Green bars, T cell GFP expression; blue bars, B-cell labeling; both signals are used to assess the significance of signal strength. Gray bars, TCR clones with signals below positive activation and/or QC thresholds. C) Left panel, PNJE3 cells expressing TCRs of interest (TT2, TT7, TT9, TT10, TT11, TT13, and TT17) specific for class II-presented epitopes within the *C. tetani* tetanospasmin protein co-cultured with DR4^+^ B-LCLs transduced with select cognate minigenes (minigene coordinates within the TetX protein listed). Right panel, epitope mapping of seven tetanus toxin (TT)-reactive TCRs following co-culture with the set of overlapping minigenes. D) Left panel, cognate validation for anti-SARS-CoV-2 TCRs. Five TCRs were co-cultured with HLA-matched allogenic B-LCLs expressing Wuhan strain/variant Spike protein minigenes. Right panel, sequence alignment for the four Spike protein variants depicting published epitopes of the respective TCRs. E) Histograms of GFP expression by CAR1 or CAR2-transgenic PNJE3 cells following co-culture with ligand expressing Du145 cells at two CAR:Du145 cell ratios (left 1:1, right 1:3).

Next, we applied the platform to epitope mapping in infectious disease. Effective vaccines contain epitopes known to elicit natural immune responses, and mapping these requires high-throughput TCR deconvolution. For this, we created a series minigenes spanning the *C. tetani* tetanospasmin (TetX) protein, the inactivated form of which is delivered as the tetanus toxoid (TT) vaccine. While several TCRs from TT-vaccinated donors and targets had already been used to benchmark the system (Figure 1), more precise epitope coordinates for each of the set of TetX-reactive TCRs were unknown. By identifying overlapping constructs which activated reporter T cells expressing a given TCR, we could map the cognate HLA-DR4 presented epitopes within the full-length protein (Figure 2C). Notably, we observed robust activation of TCRs specific for distinct epitopes within the same minigene (TetX 612-1011), demonstrating that larger constructs efficiently present multiple immunogenic epitopes, mimicking physiological antigen processing and implicitly multiplexing our deconvolution assay with respect to the number of presented epitopes generated for a given protein target.

In addition to characterizing immune responses upon vaccination to improve vaccine design strategies, the discovery of immunogenic epitopes from natural infections can guide optimal first-line vaccine design. Therefore, we validated a series of recently published^17, 18^ coronavirus spike (S) protein-reactive CD4 TCRs identified via epitope screening in SARS-CoV-2 patients. Despite the unavailability of autologous APCs or full donor haplotype information, we functionally validated several published TCRs using a panel of allogenic donor APC lines (Figure 2D, left). Furthermore, we interrogated the specificity of these TCRs against spike protein variants and observed that several of the TCRs were unresponsive to the more recently emerged Delta and Omicron variants (Figure 2D, right), indicating that T-FINDER can be used to promptly survey emerging variants of concern against existing T cell immunity, identify potential escape mutations, and guide the design of next-generation vaccines. These studies together illustrated the relevance of our system to a wide range of applications and its potential for scalability to larger numbers of TCRs and putative epitopes.

To demonstrate the flexibility and capacity of T-FINDER to support receptors beyond canonical class I/II HLA-restricted αβTCRs, we transgenically expressed a pair of chimeric antigen receptors (CAR1 and CAR2) in the PNJE3 line and co-cultured these reporter cells with the epithelial cell line Du145 exogenously expressing the CAR target. While both reporter CAR-T cells generated robust signals at a 1:3 T:Du145 ratio, a lower abundance of antigen-presenting cells (1:1) enabled behavioral discrimination between the two CARs, with CAR2 possessing greater sensitivity to the target but measurable tonic activation in its absence (Figure 2E). Based on these results, this assay can be used to assess the activity of large CAR variant libraries, selecting those which possess the desired characteristics with respect to sensitivity and response control, and eliminating potential safety concerns stemming from aberrant CAR activity.

### TCR mapping in celiac disease

Celiac disease is an immune condition resulting from T cell driven responses to dietary glutens. These plant-derived proteins, rich in glutamine and proline residues, are modified via glutamine deamidation and processed to HLA-DQ presented peptide epitopes. The plant origin, primary sequence, and post-translational modification of these epitopes presented potential challenges to T-FINDER’s endogenous minigene processing approach. To address this, we tested a series of published DQ2.5- and DQ8-restricted TCRs^19–24^ identified in celiac disease (CeD) patients. First, we generated 400 amino acid long minigene equivalents of previously identified cereal gliadin peptides (Figure 3A), substituting glutamic acid for glutamine residues at known deamidation sites. We tested a small cohort of TCRs against this minigene panel using HLA-DQ matched B-LCLs, observing significant activation signals for the expected interactions across the panel (Figure 3B). We then screened a matrix of CeD TCRs against the minigene panel in an experimenter-blinded fashion. In addition to the initial series of benchmarking TCRs, we successfully profiled 14/17 additional clonotypes with respect to their published specificities (Figure 3C), without detecting any false positive interactions. One important aspect of CeD pathophysiology is the possibility that gluten-reactive TCRs arise from cross-reactivities with gut microbiota^20^. Notably, we observed cross-reactivity to the g8pw65 bacterial epitope for one TCR which had not been previously documented.

**Figure 3.**
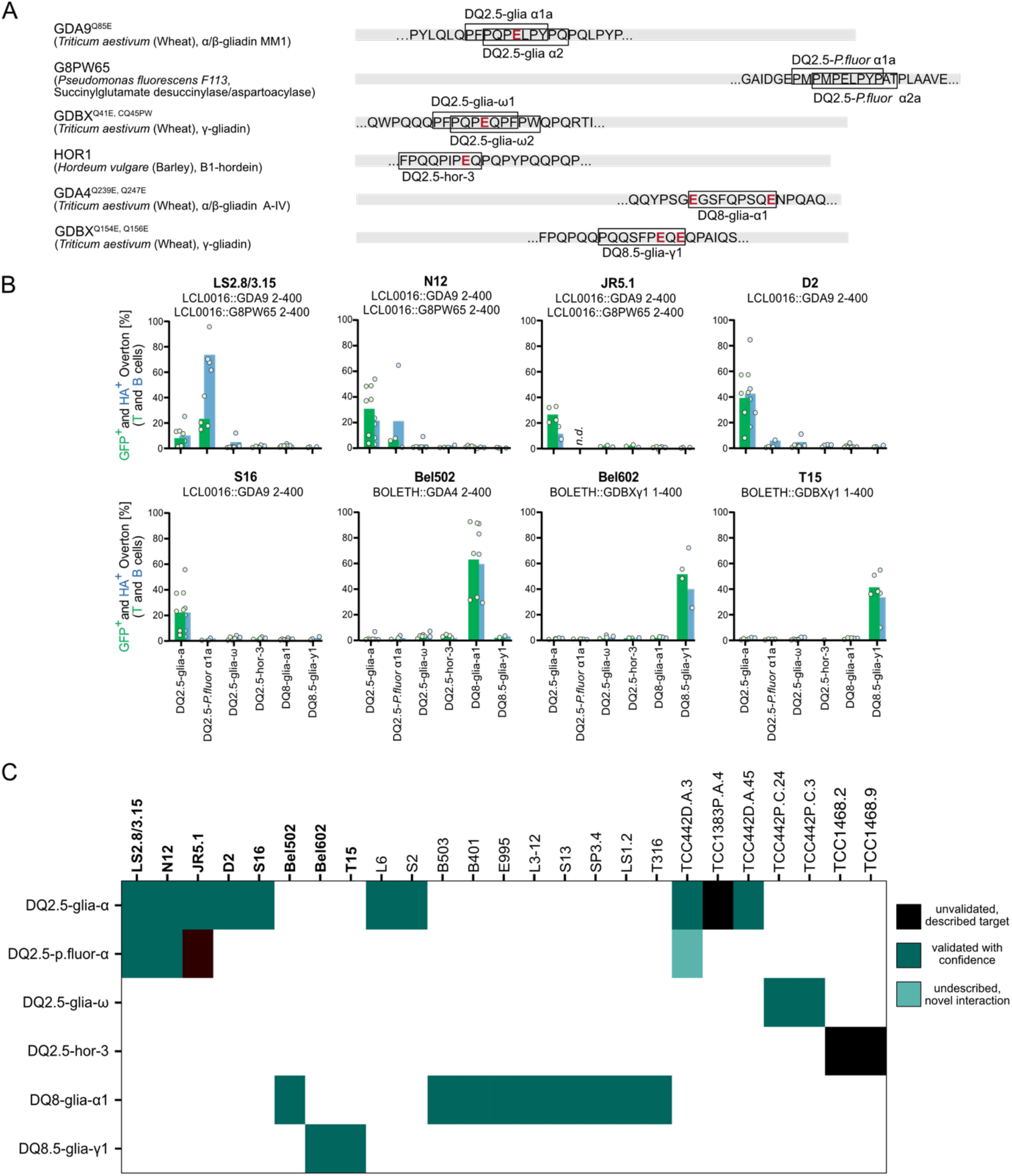
Validation of cereal gluten epitope-specific TCRs in celiac disease. A) CeD peptide epitopes (black boxes) within the 400-amino acid minigenes used in this study. Gene and species sources are listed to the left, Q-to-E mutations are indicated in red. B) Small-batch TCR testing using HLA-DQ matched allogenic B-LCLs. Example results of co-culture between the minigenes illustrated in (A) and eight CeD TCRs with published specificities (provided above each respective plot). DQ2.5-restricted TCRs in culture with LCL0016, a DQ2.5-positive B-LCL; DQ8-restricted TCRs in culture with BOLETH cells. C) Blinded screen of CeD TCRs (labeled across the top) with minigenes containing the respective CeD epitopes (labeled down the left). Black boxes indicate expected reactivities not observed in this experiment. Green boxes indicate expected (dark) and novel (light) reactivities.

### Scaling to putative target libraries for autoimmune disease

Following these pilot studies, we explored the applicability of T-FINDER to libraries on the order of 10^2^ protein targets, analogous to patient-specific tumor mutanomes, viral genomes, or tissue-specific genes representing putative autoantigens. For this, we designed a minigene library containing several autoantigens with known association to rheumatoid arthritis, as well as 162 viral and bacterial proteins associated with common infections and/or autoimmune reactions, originally intended as a counter screening tool for autoimmune disease-derived TCRs (and including TetX minigenes as positive controls). Based on our observations that reporter T cells react to cognate-presenting cells even when diluted by non-targets (Supplemental Figure 4A), we designed an arrayed antigen multiplexing system which would allow us to definitively identify the correct target. To establish a given array, putative antigen minigenes are distributed across a series of wells according to a set-generating algorithm. Key to this design is the constraint that no two putative targets co-occur in more than one well of the library array; in this way, the containing activated T cells serve as a fingerprint which is resistant to both false positive and false negative signals (Figure 4A). This indexed array approach removes the need for downstream deconvolution or target sequencing while still compressing screen size approximately five-fold.

**Figure 4.**
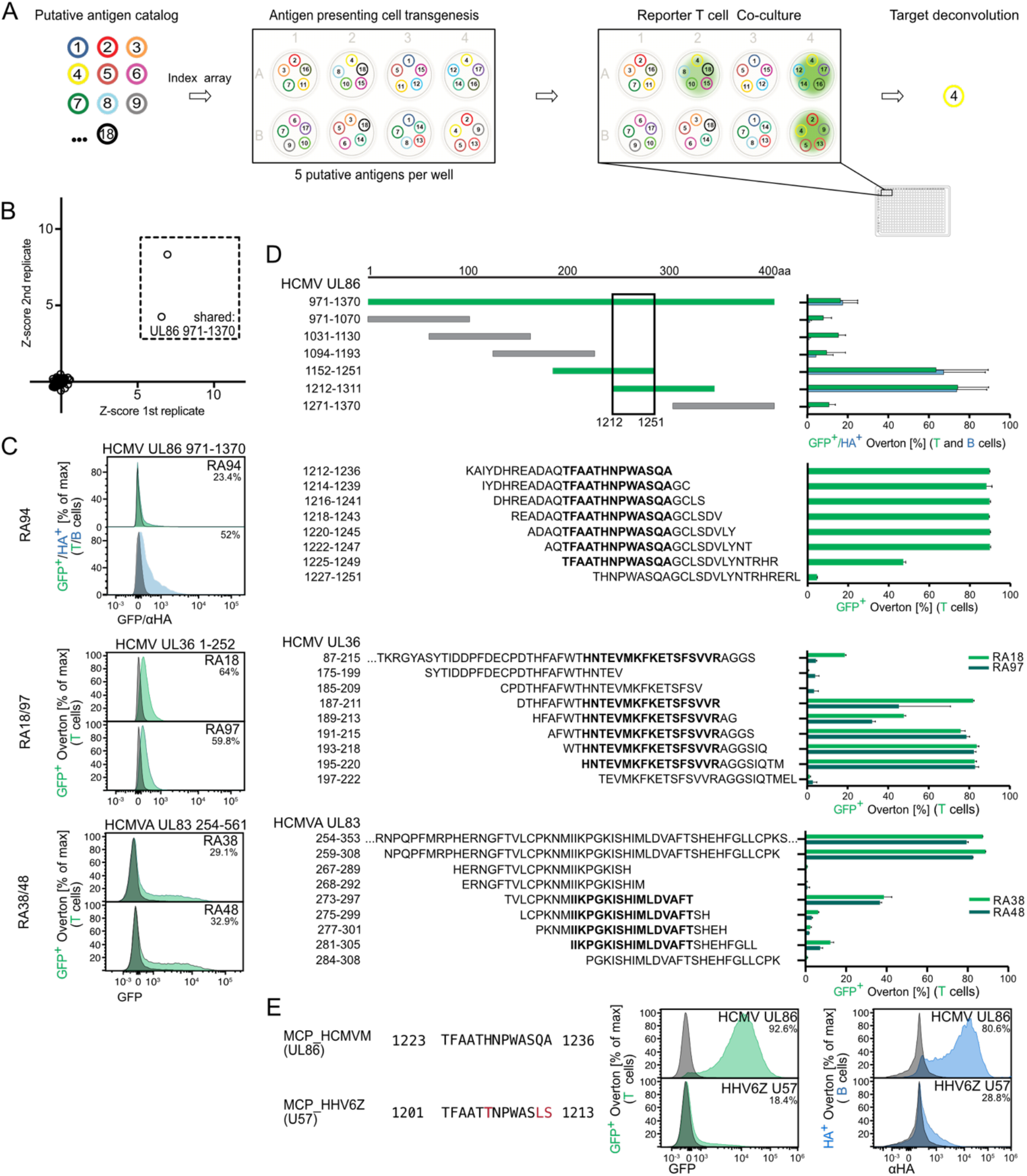
Epitope discovery, mapping, and cross-reactivity for rheumatoid arthritis TCRs. A) Schematic depiction of the algorithmic antigen array principle: putative targets (numbered 1-18 in this example) are assigned to a well (8 wells in this example) such that no two co-occupy a well more than once, while each target appears multiple times in the plate. Target pools are transduced into B-LCLs and reporter T cells are added for co-culture. Microscopic and/or flow cytometric analysis identifies positive wells, the combination of which immediately indicates the cognate minigene. B) Identification of a fragment of the HCMV UL86 protein as a functional ligand of RA94, a rheumatoid arthritis patient derived TCR through multiplexed minigenes as well as individual validation. D) Medium-(top plot) and high-(bottom three plots) resolution minigene scanning for epitope-containing regions within cognate 400 amino acid minigenes. Bold sequences indicate the minimal epitopes of the respective TCRs in the adjacent plots from panel (C). E) Cross-reactivity between the RA94 mapped minimal epitope, provided as an *in vitro* loaded peptide (upper panels), and homologous peptide from a genomically integrated herpesvirus (lower panels).

To assess the target identification capacity of this approach, we tested 11 TCRs representing enriched clonotypes from inflamed joints of HLA-DR^+^ rheumatoid arthritis patients^25^, the specificities of which were initially blinded to the experimenters. Screening each TCR against the array in duplicate, we confirmed the positive control TetX TCR targets (Supplemental Figure 4B) and identified multiple positive wells sharing a single target for three TCRs (RA94, Figure 4B, and RA18/RA97, Supplemental Figure 4C). After the initial arrayed screen, we validated each of the targets in single-minigene co-culture (Figure 4C, Supplemental Figure 4D). All three successfully deconvoluted TCRs recognized herpesviral targets, with RA18 and RA97 both reacting to the UL36 tegument protein of herpes simplex virus1 and RA94 reacting to the UL86 major capsid protein of the human cytomegalovirus. Notably, these TCRs were identified in three individual donors, suggesting a possible link between antiviral immune responses and RA etiopathology.

### Minimal TCR epitopes can rapidly be identified via successively smaller antigenic fragments

One advantage of using larger minigene-encoded targets is the ability of APCs to process and present multiple epitopes from a single minigene (see for example the responses of TT9, TT10, and TT11 to tetanospasmin fragments in Figure 2C). This enables our platform to cover a much broader epitope space (for example the full human proteome) with a smaller number of library elements while ensuring adequate screening depth. For many applications, however, it is relevant to identify cognate peptides at higher resolution. Libraries of overlapping peptides are one option, but cannot always be synthesized in a timely or efficient manner. As a fast and flexible alternative, we applied our minigene approach to identify minimal TCR epitopes over, at most, two successive rounds of construct design. In this way, we were able to map the minimal epitopes of the previously deconvoluted RA-derived TCRs, as well as those of two additional RA-derived TCRs (RA38 and RA48) whose targets did not initially pass QC thresholds in the arrayed screen but could be validated in single-minigene co-culture (Figure 4D). This final step demonstrates the general utility of our TCR deconvolution system, beginning with a clonotype of interest and unknown ligand through to a detailed mapping of the cognate peptide:HLA complex.

Amino-acid epitope resolution allows us to revisit the concept of epitope mimicry in autoimmune disease, in this case by enabling a more defined homology search between viral targets and potentially cross-reactive human proteins. In principle, no TCR has a single peptide epitope; rather, TCR specificities cover an “epitope space” with a distinct affinity for each member^26^. In fact, this space can widely accommodate a variety of functional epitopes, even if the peptide sequences are not highly homologous^27^. T-FINDER’s ability to quickly progress from *de novo* ligand identification at low resolution (400 amino acids) to minimal epitope mapping enables rapid exploration of this multispecificity. As an example, a BLAST homology search of the minimal epitope for RA94 against the human reference proteome identified a similar peptide which is encoded by a genomically integrated viral capsid protein^28^. Indeed, this fragment also activates RA94-transgenic reporter cells (Figure 4E). Continuing work mapping such epitope spaces will support the connection between the presence of apparently antiviral TCR clonotypes in autoimmune tissues and their role in contributing to inflammation via additional, self-derived targets.

## Discussion

Here we describe the development and validation of T-FINDER, a flexible TCR deconvolution platform. Key features of T-FINDER include a highly sensitive promoter capable of transducing signals from TCRs to CARs, and likely related receptors (e.g., γδ T cell, NK) that utilize the same signaling pathway. The inducible cassette, engineered into the PNJE3 reporter clone, provides for labeling of activated reporter cells and the activating B-LCL presenting cell. By engineering the reporter line, rather than the presenting cell, T-FINDER remains compatible with a variety of antigen presentation options including different cell types, HLA classes, and non-HLA ligands – as evidenced by our experiments with CAR constructs. Notably, T-FINDER interrogates an individual’s entire HLA repertoire, given the ability to use donor derived B-LCLs as APCs – a unique capability among deconvolution approaches.

The high-throughput capacity of T-FINDER is an important feature due, in part, to the use of 400 amino acid long minigene constructs to encode the antigen library. Typical minigene libraires consist of shorter (<100 aa) fragments due to inefficient processing of minigene encoded targets by class II HLA. However, T-FINDER overcomes this limitation by use of a PES upstream of the antigen (Pinamonti et al, manuscript in preparation). As demonstrated by the TetX experiments, PES-enabled minigenes efficiently present multiple distinct epitopes encoded within 400-aa constructs. In addition to endogenous antigen processing, T-FINDER can accommodate post-transcriptional modifications (PTM) with minor adaptations (e.g. genetic introduction of deamidated residues or the introduction of modifying enzymes into the B-LCL). Hence, T-FINDER’s antigen libraries require fewer distinct constructs to provide greater antigen breadth and depth relative to other approaches.

The importance of sensitive cellular reporters of T cell activation has been highlighted by many recently described studies establishing such systems^29–31^. While some of these systems’ main goal is to dissect individual TCR signaling pathways or focus on fast proximal signals such as calcium flux via NFAT, we have integrated all activation-related activity onto a single, compact locus control region. At the same time, we observe little to no aberrantly induced signal in the absence of TCR-cognate epitopes, indicating that, while highly sensitive, our promoter is not susceptible to false positive activation events or tonic TCR signals.

While the potently inducible promoter element of our reporter cassette can be re-engineered for numerous applications beyond the current Jurkat-based line (including murine T cell lines and human primary T cells), the PNJE3 clone can be easily adapted to any of several antigen-presenting cell possibilities. If primary donor material is available, reporter T cells can be co-cultured with peripheral blood mononuclear cells, tissue explants, or organoids to test for functional reactivity. Instead of using B-LCLs as APCs, tumor cell lines (with or without donor-derived neo-epitope libraries) may be advantageous for observing physiological T cell responses in an immuno-oncology setting. Finally, allogenic or engineered APCs can be used to confirm HLA restriction if TCR identification was initially performed in an unbiased manner.

Computational approaches to identify TCR ligands are abundant, with many able to accurately predict cognate TCR sequences for a known epitope^32^. However, identifying novel ligands for a given TCR sequence presents a much greater *in silico* challenge, and the accuracy of these models is limited by the trustworthiness of their training sets: many publicly available ligandome sets are either based on TCRβ chain sequencing only (rather than complete αβ pairs) and/or have not been functionally validated. In fact, early attempts to use more than 15 class II TCR:autoantigen pairs obtained from public databases as positive controls for our system failed to activate the PNJE3 line, either as minigenes or peptides (data not shown). In contrast, T-FINDER deconvolutes and functionally validates TCR:ligand pairs, and can also be used to simultaneously increase the size and quality of training data sets for prediction models.

Isolating TCRs from their primary T cells for insertion into a reporter cell line may initially seem extraneous. However, measuring the activation status of primary T cells presents challenges with respect to overall viability and the ability to activate tolerized or exhausted cells, which may lead to the loss of relevant clonal sequences. Some applications in TCR and ligand discovery will still require an enrichment step for ligand (set)-reactive TCRs, such as when donor samples are only available from the peripheral blood, which particularly for CD4 T cells has very complex TCR repertoires^33^. However, when fewer enriched clones are expected, for example from tumor-infiltrating lymphocytes or inflamed autoimmune tissue, it will be possible to sequence TCRs and move directly to target identification, removing a potentially bias-introducing step.

An important feature of using genetically encoded TCR ligands rather than directly loaded peptides is their usage of cellular antigen-processing machinery. In effect, we increase physiological relevance by offering antigen-presenting cells larger protein moieties, which must be naturally cleaved and loaded onto HLA by the APC. This approach can potentially identify epitopes not predicted based on previous, mass spectrometric ligandome studies^34^, which may not comprise a complete catalog of peptides that can be presented by a given HLA molecule. Furthermore, physiological antigen processing allows APCs to dictate peptide lengths rather than be constrained by synthetic peptide design, which is particularly advantageous for extended, variable-length class II-presented epitopes^35^. Synthesized peptides or libraries thereof cannot be feasibly scaled without an *a priori* estimate of epitope length, potentially resulting in false negative interactions with TCRs that require extended residues. It is more efficient and less biased to assign the task of antigen processing to the APCs themselves. Using minigenes, a reasonably small epitope can be identified, followed by the minimal functional sequence required for peptide processing and presentation. This approach can also characterize residues affecting antigen presentation that are not directly involved in TCR binding, but rather in the ability of APCs to process the antigens, which opens a new avenue to manipulate T cell epitopes to increase or decrease their immunogenicity, depending on the desired outcome.

With several alternatives to TCR and epitope discovery currently available, but none satisfying our strict requirements for both potency and flexibility, we have generated a multicomponent system which can be adapted to serve many needs in T cell based applications. More importantly, T-FINDER drives new biological insights into immune cell specificity at a pace that was previously not possible, enabling and improving the development of precision diagnostics and therapeutics.

## Acknowledgements

The authors would like to thank Jeroen Van Houdt, Viggo Van Tendeloo, Nina Papavasiliou, and Murray McKinnon for their support, input, and guidance throughout the project. We would like to express our gratitude to all donors and clinical units for providing and collecting samples used in the study. This work was supported by BioMed X funding to JML from the Janssen Pharmaceutical Companies of Johnson and Johnson.

---

**Supplemental Figure 1.**
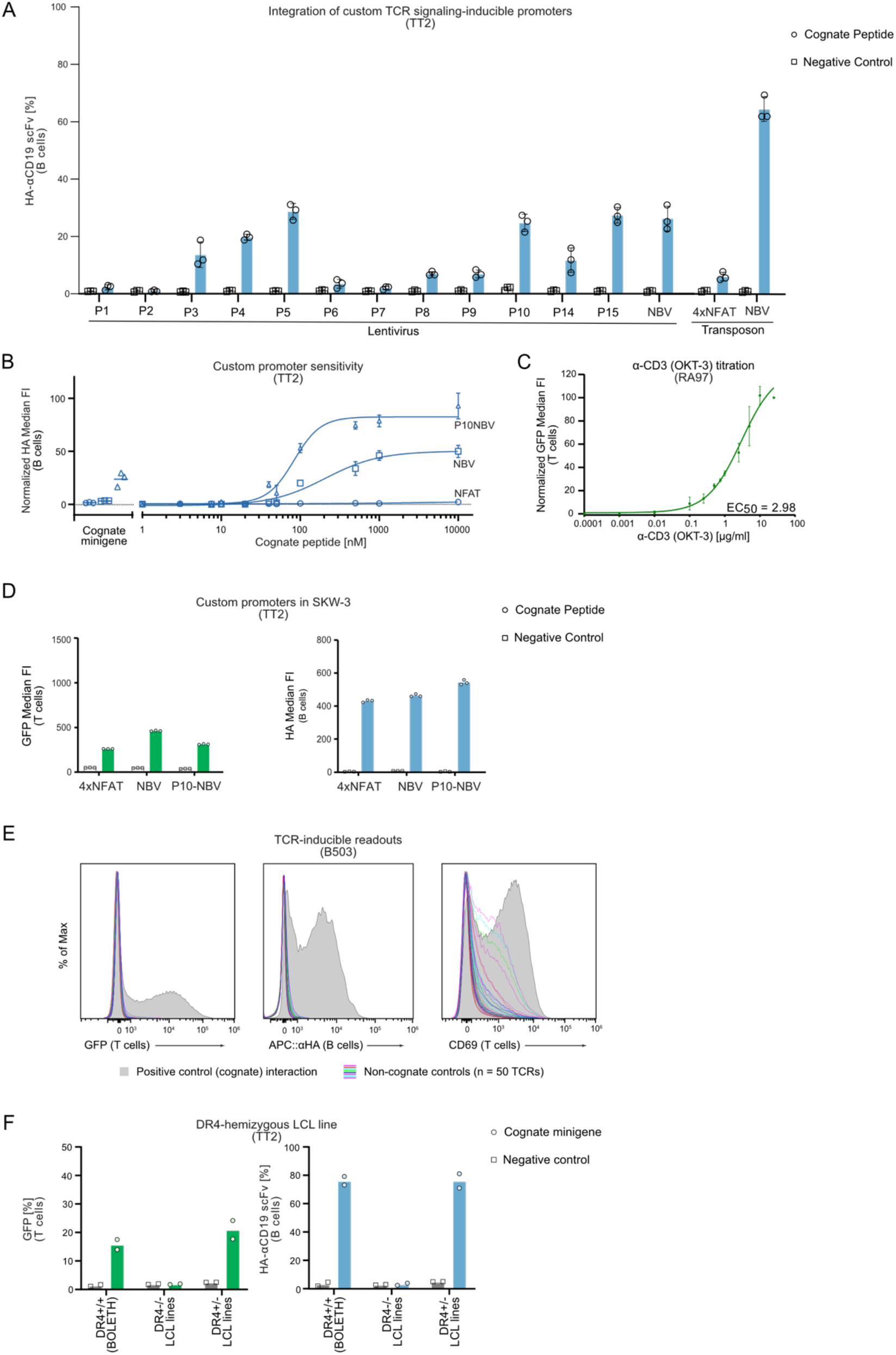
TCR activation reporters and minigene antigen constructs. A) Labeling of cognate B-LCLs (via αCD19 scFv secreted by activated reporter T cells) for a series of bespoke TCR signaling-inducible promoters. Promoters with strongest activity were stably integrated as transposons (rightmost bars), together with a 4xNFAT binding sequence promoter for comparison. B) Minigene and peptide dilution curves (HA-tagged αCD19 scFv) of a cognate epitope for three transposon-integrated inducible reporter cassettes (bulk T cell population prior to single-clone selection). C) Titration curve for TCR-independent reporter T cell activation of one rheumatoid arthritis-derived TCR using an αCD3 antibody (OKT-3). D) Inducible promoter activity (lentiviral integration) in SKW-3 cells. E) T-FINDER readouts (left, T cell GFP expression; middle, B-cell αCD19 labeling) of reporter T cell activation (cognate minigene co-culture, solid histograms) and background noise (negative control TCRs with known targets not expressed by the APCs, colored lines), relative to T cell CD69 surface expression as a readout (right). F) Bar graphs showing TT2-transgenic reporter T cell activation (left, GFP expression; right, B-cell labeling) following co-culture with a homozygous HLA-DR matched cell line (BOLETH, DR4+/+), a DR4-negative B-LCL, and a hemizygous (DR4+/-) HLA-haplotyped LCL line pulsed with cognate peptide.

**Supplemental Figure 2.**
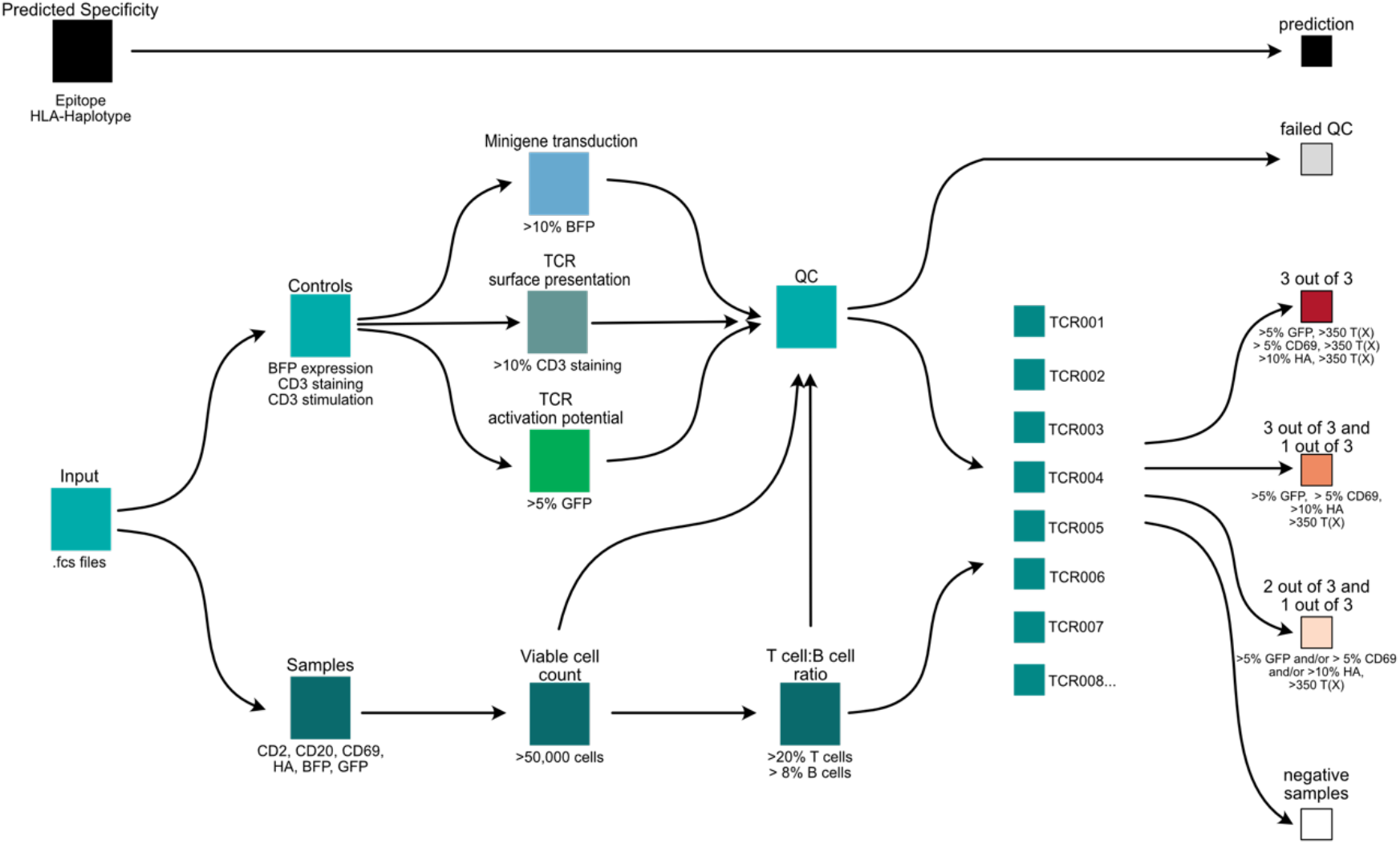
**Analysis pipeline for celiac disease TCRs**, cf. Figure 3C. This workflow depicts the parameters analyzed to ensure robust and high-quality validation calls on large, blinded TCR:target screens. The following experimental parameters are considered: successful TCR expression by reporter cells (>10% of CD3-positive T cells), activation potential upon ligand-independent αCD3 stimulation (>5% of GFP-positive T cells), successful minigene expression by B-LCLs (>10% of BFP-positive B cells), and acceptable T:B ratios in the co-culture. Only samples that pass all determined QC thresholds are eligible for analysis. Upon passing QC, GFP and CD69 (T cells) and αHA (B cell labeling) are used to determine TCR activation signal strengths and call confidence.

**Supplemental Figure 3.**
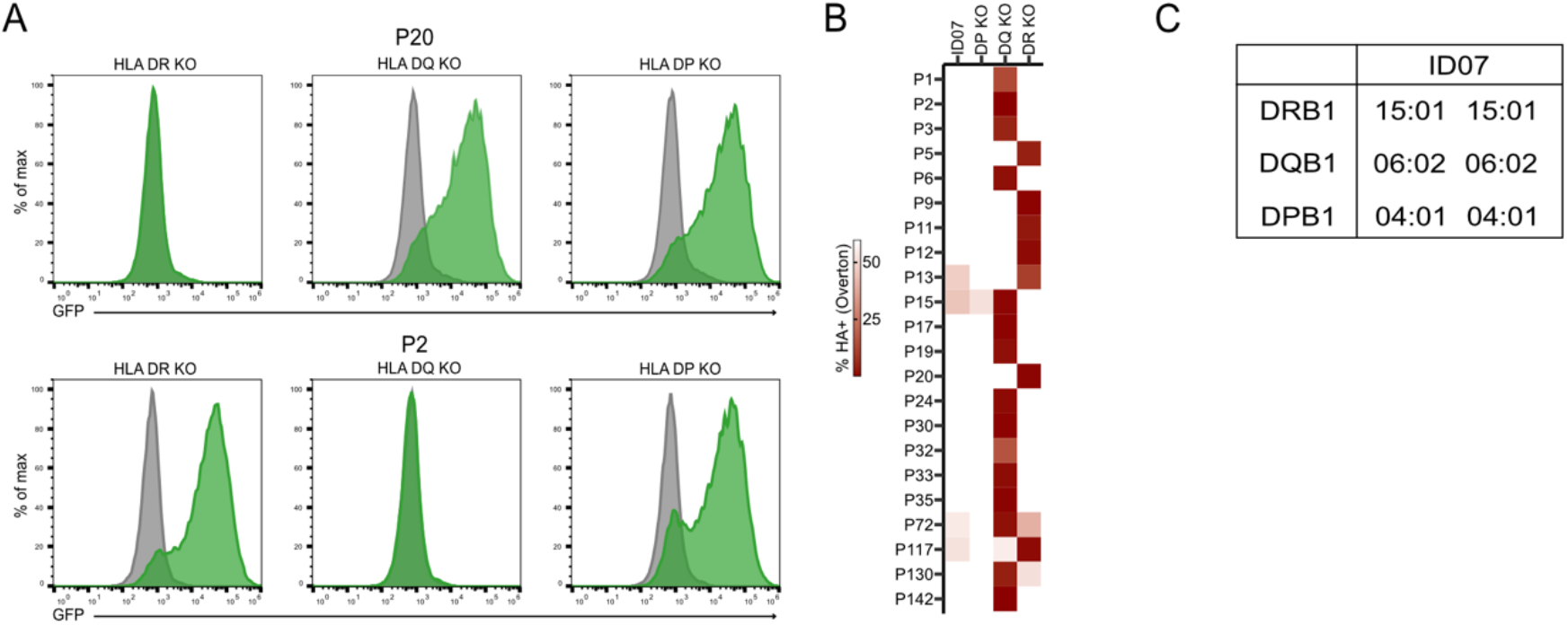
HLA restriction mapping of glioma peptide-reactive TCRs. A) Example flow cytometry data for two TCRs (P20, upper panels; P2, lower panels) co-cultured with class II HLA-deficient autologous B-LCLs. Gray histograms, control peptide-loaded APCs; green histograms, cognate peptide-loaded APCs. TCR specificity for the cognate epitope in complex with the restricted HLA allele is lost upon deficiency of the respective allele (e.g. HLA-DR for P20, HLA-DQ for P2). B) TCR restriction analysis for the full set of deconvoluted neo-epitope specific TCRs from patient ID07. C) Patient ID07 class II HLA haplotype.

**Supplemental Figure 4.**
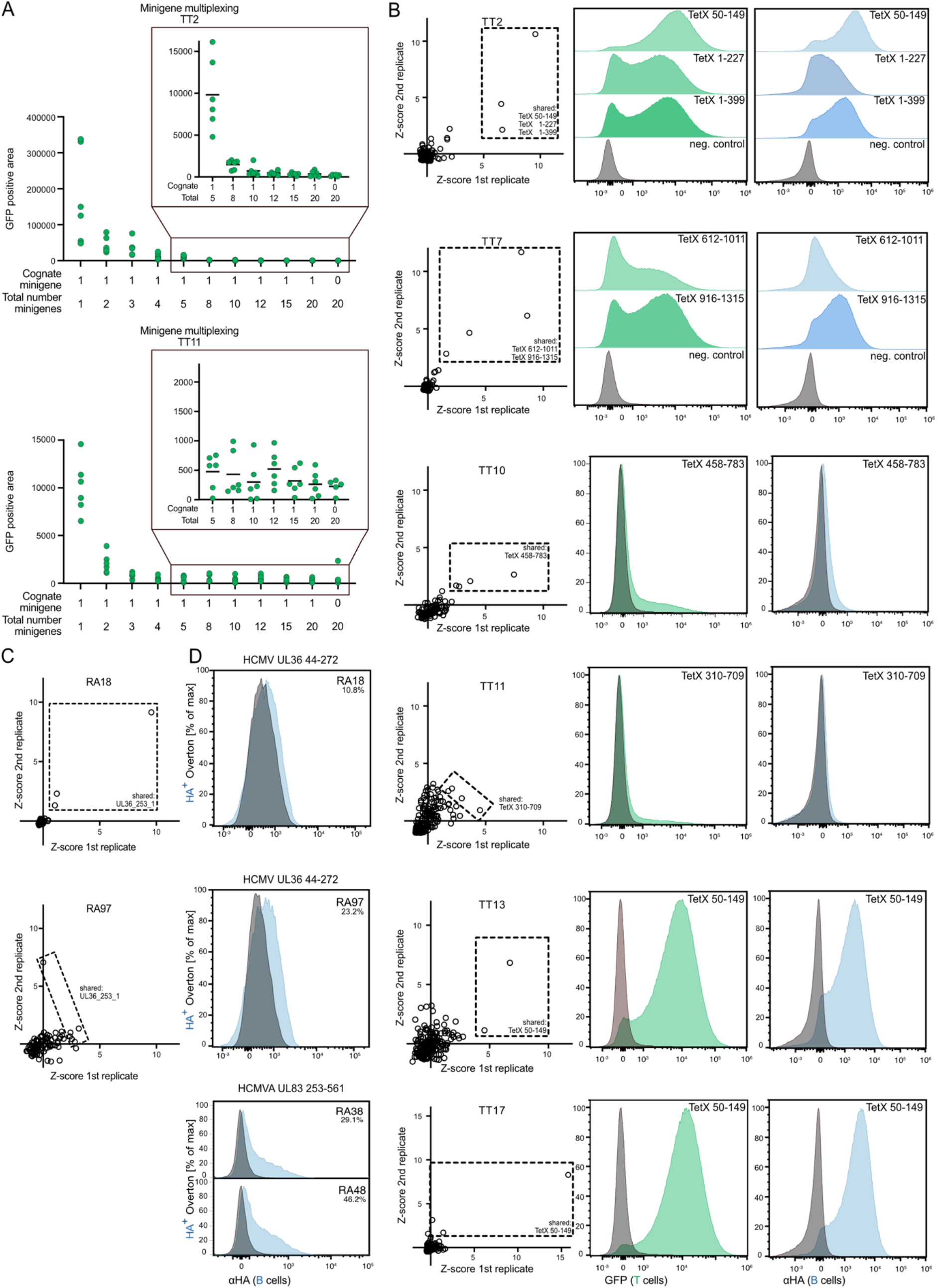
TCR ligand identification from putative target libraries. A) Putative target multiplexing. An increasing number of non-cognate targets was added at equimolar ratios to the mixture of minigenes used to transduce B-LCLs prior to co-culture with reporter T cells bearing TCRs (the strongly reactive TT2 or the weakly reactive TT11) specific for a single target in the mixture. B) Plate-based antigen discovery for six tetanus toxoid-reactive TCRs. Identification of positive wells with shared minigenes (dashed boxes) in an array containing 162 putative targets across 96 wells. The screen was performed in duplicate (x and y axes) to better identify reproducibly positive wells. Histograms to the right: reporter T cell activation in single-target minigene co-culture for the respective arrayed screen results on the left. C) Plate-based antigen discovery for two and D) single-target validation of four rheumatoid arthritis TCRs.

## Notes

### Competing Interest Statement

JML, NF, LF, YH, JZ, KK, VP, and MC are listed as inventors on a patent application describing a System and Methods for Identifying T cell Receptor Ligands. JML, TL, and TS are employees of BioMed X GmbH; VD, LF, YH, JZ, KK, AMF, VP, and MC were employees while conducting work related to this study. MP and EWG are founders of Tcelltech GmbH. NF, FS, CVH, HW, BVS, and AK are/were employees of Janssen while conducting work related to this study.

